# Lysyl oxidase-dependent subendothelial matrix stiffening promotes RAGE-mediated retinal endothelial activation in diabetes

**DOI:** 10.1101/2022.08.31.505952

**Authors:** Sathishkumar Chandrakumar, Irene Santiago Tierno, Mahesh Agarwal, Nikolaos Matisioudis, Timothy S. Kern, Kaustabh Ghosh

## Abstract

Endothelial cell (EC) activation is a crucial determinant of retinal vascular inflammation associated with diabetic retinopathy (DR), a major microvascular complication of diabetes. We previously showed that, similar to abnormal biochemical factors, aberrant mechanical cues in the form of lysyl oxidase (LOX)-dependent subendothelial matrix stiffening also contribute significantly to retinal EC activation in diabetes. Yet, how LOX is itself regulated and precisely how it mechanically controls retinal EC activation in diabetes is poorly understood. Here we show that high glucose-induced LOX upregulation in human retinal ECs (HRECs) is mediated by proinflammatory RAGE (receptor for advanced glycation end products/AGEs). HRECs treated with methylglyoxal (MGO), an active precursor to the AGE MG-H1, exhibited LOX upregulation that was blocked by a RAGE inhibitor, thus confirming the ability of RAGE to promote LOX expression. Crucially, as a downstream effector of RAGE, LOX was found to mediate both the proinflammatory and matrix remodeling effects of MGO/RAGE, primarily through its ability to crosslink/stiffen matrix. Finally, using decellularized HREC-derived matrices and a mouse model of diabetes, we demonstrate that LOX-dependent matrix stiffening feeds back to enhance RAGE, thereby achieving its autoregulation and proinflammatory effects. These fresh insights into the regulation and proinflammatory role of LOX-dependent mechanical cues may help identify new therapeutic targets to block AGE/RAGE signaling in DR.

## Introduction

Diabetic retinopathy (DR) is a common microvascular complication of diabetes that remains the leading cause of preventable vison loss in the working-age population (1,2). In the early stages of DR, hyperglycemia leads to retinal inflammation that activates retinal vascular endothelial cells (ECs). Activated retinal ECs, in turn, upregulate intercellular adhesion molecule-1 (ICAM-1), a crucial proinflammatory EC adhesion molecule that binds circulating leukocytes and, thereby, contributes to the development of early vascular lesions of DR (viz. vascular atrophy and hyperpermeability) (3,4). However, precisely how retinal ECs become activated and express ICAM-1 in diabetes remains insufficiently understood.

Recent findings have revealed that, under high glucose (HG) conditions, retinal ECs increase the expression of lysyl oxidase (LOX) (5,6), a copper-dependent amine oxidase that crosslinks and stiffens extracellular matrix (5,7). These findings correlate with LOX upregulation observed in the retina and vitreous of diabetic animals and humans, respectively (5,6,8). Crucially, our studies showed that LOX-mediated subendothelial matrix stiffening alone is sufficient to promote retinal EC activation (ICAM-1 expression) in vitro (5), a finding that aligns with the proinflammatory effects of matrix stiffening on cardiovascular and lung ECs in atherosclerosis and sepsis, respectively (7,9). This ability of subendothelial matrix stiffening to control EC behavior is attributed to endothelial mechanotransduction, a process by which ECs transduce matrix-based mechanical, structural, and adhesive cues into intracellular biochemical signals that regulate cell behavior at both translational and transcriptional levels (5,10-12). Given this newly identified role of LOX in the mechanical control of retinal EC activation, it becomes important to elucidate both the regulation and proinflammatory role of retinal LOX in diabetes.

LOX is known to be upregulated in inflammatory conditions such as sepsis and atherosclerosis (7,13). Since HG also promotes proinflammatory signaling by upregulating RAGE (receptor for advanced glycation end products), we asked whether HG-induced LOX upregulation in retinal ECs is mediated by RAGE. RAGE is the chief receptor for advanced glycation end products (AGEs), which are formed by nonenzymatic glycation of proteins and lipids in diabetes and aging. Notably, the proinflammatory AGE/RAGE signaling is significantly enhanced in the retinas of diabetic subjects where it contributes to retinal vascular inflammation and degeneration by inducing sustained NF-κB-dependent ICAM-1 upregulation, leukostasis, breakdown of blood-retinal barrier (BRB), and EC/pericyte apoptosis (14-16). Despite their important implications in DR pathogenesis, direct inhibition of AGE or RAGE has proven challenging in clinical trials due to drug safety issues and low efficacy of the drug (17-19). Thus, determining the role of AGE/RAGE signaling in retinal endothelial LOX expression may not only provide mechanistic insights into LOX regulation in diabetes but, crucially, also implicate LOX as an alternative (downstream) target to block AGE/RAGE signaling.

Here we report that HG and methylglyoxal (MGO), precursors of AGE formation and subsequent RAGE activation, increase LOX expression and activity in retinal EC cultures in a RAGE-dependent manner. Importantly, inhibiting LOX alone blocked the ability of MGO/RAGE to cause EC activation (ICAM-1 expression and monocyte-EC adhesion) and subendothelial matrix remodeling (increased matrix stiffness and alignment). Finally, using our mouse model of diabetes and retinal EC cultures, we demonstrate that LOX feeds back to increase RAGE expression, thus providing a potential mechanism for both LOX autoregulation and its observed proinflammatory effects (5).

### Research Design and Methods

#### Experimental Animals

All animal procedures were performed in accordance with the Association for Research in Vision and Ophthalmology (ARVO) Statement for the Use of Animals in Ophthalmic and Vision Research and approved by University of California Riverside Institutional Animal Care and Use Committee. Diabetes was induced in adult (8 wk-old) male C57BL/6J mice (Jackson Laboratory, Bar Harbor, ME, USA) using our previously reported protocol (5) that is described in *Supplemental Material*. Diabetic mice were treated with a specific and irreversible LOX inhibitor β-aminopropionitrile (20) (BAPN; 3 mg/kg through drinking water; Catalog no. A3134-5G; Sigma-Aldrich, St. Louis, MO, USA) for 20 weeks prior to euthanasia and collection of eyes for further analyses.

#### Cell culture and treatments

Human retinal endothelial cells (HRECs) were purchased (catalog no. ACBRI 181; Cell Systems corp., Kirkland, WA, USA) and cultured as previously reported by us (5). Human THP-1 monocytes were purchased from ATCC (Manassas, VA, USA) and cultured in vendor recommended medium as described in *Supplemental Material. <u>In vitro treatment regimen</u>*: Confluent HRECs (passages 5-8) grown in regular culture medium containing 5.5 mM glucose were either left untreated (UT) or treated with methylglyoxal (MGO) ± 0.1 mM BAPN or 0.1 μM RAGE Antagonist FPS-ZM1 (Calbiochem-Sigma Millipore) for 10 days prior to use in the different assays. All cultures were supplemented with 200 μg/mL L-ascorbic acid (Catalog no. A4034; Sigma-Aldrich) to facilitate matrix deposition. To assess the role of NF-κB in MGO-induced LOX regulation, HRECs were co-treated with MGO and NF-κB inhibitor BAY 11-7082 (1 μM; Enzo Life Sciences Inc., NY, USA) for 3 days prior to assessment of LOX mRNA expression.

#### RT-qPCR

mRNA expression was assessed using our standard RT-qPCR protocol (21) that is described in *Supplemental Material*. The following gene/species-specific TaqMan primers were used for RAGE (Hs00542584_g1), LOX (Hs00942480_m1), ICAM-1 (Hs00164932_m1), and housekeeping gene GAPDH (Hs02786624_g1).

#### Western blot

Expression levels of target proteins were assessed using our standard Western blotting protocol (21) that is described in *Supplemental Material*. The following protein-specific primary antibodies were used for RAGE (catalog no. ab3611; Abcam), MG-H1 (catalog no. STA-011; Cell Biolabs, Inc., San Diego, CA, USA), LOX (catalog no. NB110-59729; Novus Biologicals, San Diego, CA, USA), ICAM-1 (catalog no. SC-8439; Santa Cruz Biotechnology, Santa Cruz, CA, USA), and loading control GAPDH (catalog no. G9545; Sigma-Aldrich).

#### ELISA

Levels of methylglyoxal adduct Methyl-glyoxal-hydro-imidazolone (MG-H1) were measured in retinal and cell culture lysates using a competitive enzyme-linked immunosorbent assay (ELISA) (Catalog no. ab238543; Abcam, Cambridge, MA, USA) as per the manufacturer protocol.

#### Monocyte-EC adhesion assay

Monocyte-EC adhesion assay was performed and ICAM-1 clustering index quantified as per our previously reported protocols (5,12) that are described in *Supplemental Material*.

#### LOX Activity assay

LOX activity was measured from HREC culture supernatant using our previously reported protocol (5) that is described in *Supplemental Material*.

#### siRNA transfection

HRECs (passage 5-6) were transfected with LOX siRNA (Catalog no. hs.Ri.LOX.13, Integrated DNA Technologies, Inc, IA, USA) employing Lipofectamine RNAiMax transfection reagent (Catalog no. 13778-030, Invitrogen, Carlsbad, CA, USA), as per the manufacturer’s protocol. Nontargeting scrambled siRNA was used as a control and transfection efficiency was evaluated by quantifying LOX mRNA.

#### Subendothelial matrix

Decellularized subendothelial matrices were obtained from HREC cultures grown in NG or MGO ± BAPN medium using our previously reported protocols (5,22) that are described in *Supplemental Material*.

#### Matrix stiffness

A NanoWizard® 4 XP BioScience atomic force microscope (Bruker Nanotechnologies, Santa Barbara, CA), fitted with a pre-calibrated PFQNM-LC-A-CAL probe (spring constant 0.075 N/m) containing a 70 nm-radius hemispherical tip (Bruker AFM Probes, CA, USA) and coupled with a Zeiss Axiovert phase contrast microscope, was used for stiffness measurement and topographical scanning of the subendothelial matrix. Stiffness was measured in the *contact mode force spectroscopy* mode by applying a 50 pN (set point) indentation force while topography of a 30 μm × 30 μm area was scanned in the *QI™ (Quantitative Imaging)* mode at a 128×128 pixel resolution using a 200 pN (set point) indentation force. Multiple force curves (n≥40/condition for stiffness and n≥1000/condition for topography) were analyzed using JPK Data Processing Software.

#### Synthetic matrix fabrication

Polyacrylamide-based synthetic matrices of tunable stiffness have been widely used by us and others as cell culture substrates to study cellular response to the mechanical properties of extracellular matrix (5,11,12). The synthetic matrices used in the current study were fabricated as per these published protocols, which are described in *Supplemental Material*.

#### Statistics

All data were analyzed using GraphPad Prism 6.01 (GraphPad Software, San Diego, CA). Statistical differences between three or more groups were assessed using a one-way analysis of variance (ANOVA) followed by Tukey’s or Dunnett’s *post hoc* multiple comparisons test, depending on whether the data followed normal distribution or not. Two-tailed unpaired Student’s *t* test was used for studies comparing two experimental groups. Results were considered significant if *p*<0.05.

## Results

### Diabetes-induced retinal EC activation is associated with RAGE and LOX upregulation

We have previously shown that diabetes and HG increase LOX expression in the retina and HRECs, respectively, that, in turn, promotes EC activation (5). To test the hypothesis that this LOX upregulation in retinal ECs is mediated by AGE/RAGE, we first assessed the degree to which HG increases AGE/RAGE levels in these cells. As shown in **Fig. 1A**, HG (but not mannitol, osmotic control) treatment of human retinal EC (HREC) cultures caused a 1.5-fold (*p*<0.001) increase in RAGE expression, which was consistent with an ∼1.5-fold increase (*p*<0.05) in retinal RAGE levels in diabetic mice **(Fig. 1B)**. Notably, this HG-induced RAGE upregulation was associated with a 1.3-fold increase in the formation of methylglyoxal hydroimidazolone1 (MG-H1), an AGE that is implicated in DR pathogenesis and activates RAGE (23), in both HREC cultures (*p*<0.01; **Fig. 1C**) and retinas of diabetic mice (*p*<0.05; **Fig. 1D**). Further, consistent with the proinflammatory effects of AGE/RAGE interactions, we observed a concomitant increase in ICAM-1 expression in both HG-treated HRECs (∼2-fold; *p*<0.001; **Fig. 1E**) and retinas of diabetic mice (∼1.7-fold; *p*<0.01; **Fig. 1F**).

**Figure 1:**
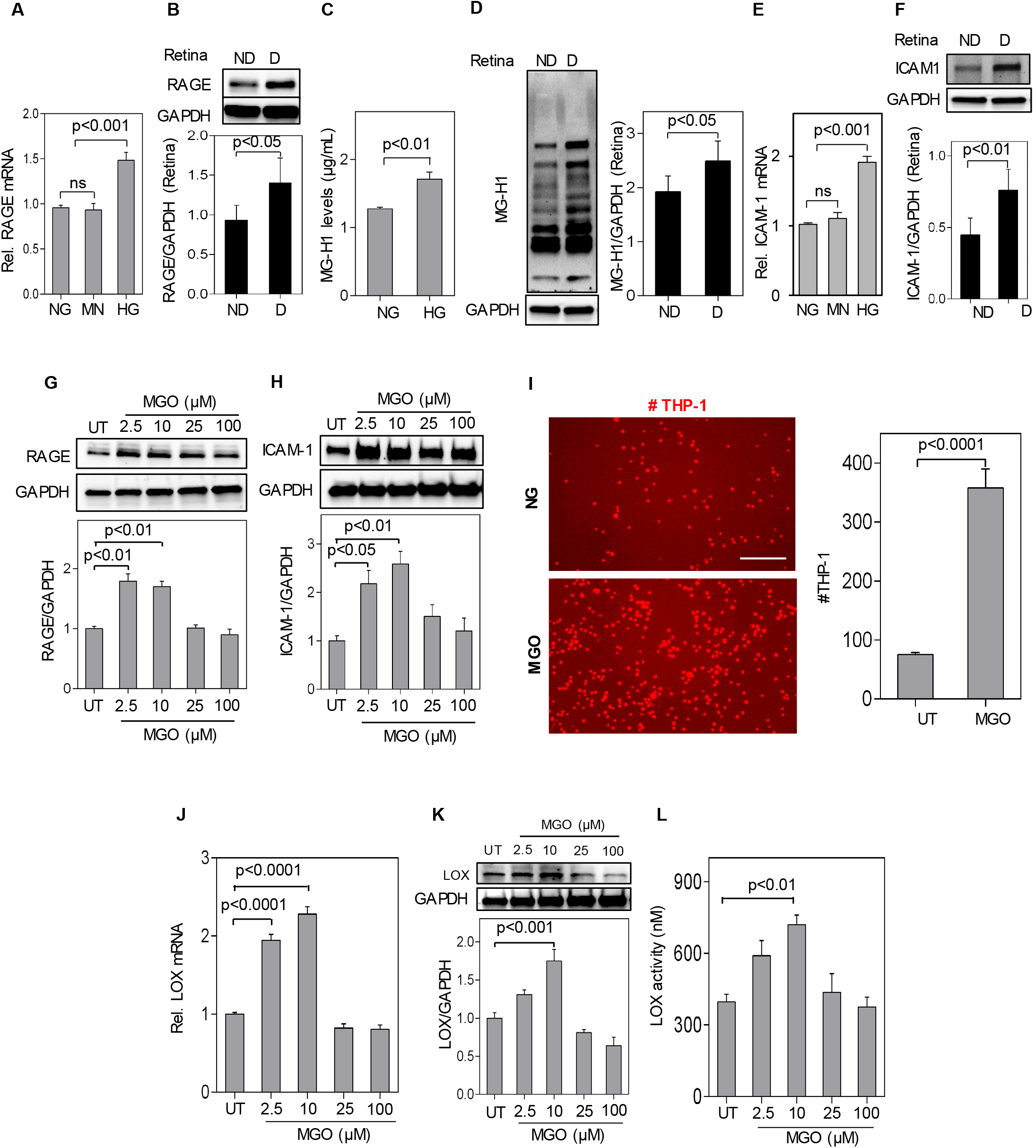
Diabetes-induced retinal EC activation is associated with RAGE and LOX upregulation. **(A)** RT-qPCR analysis of HRECs treated with normal glucose (NG), Mannitol (MN, osmotic control) or high glucose (HG) for 10d revealed a significant (1.5-fold; *p*<0.001) increase in RAGE mRNA levels under HG conditions. **(B)** Representative Western blot bands and cumulative densitometric analysis of whole mouse retinas (n=6/group) revealed significantly higher (*p<*0.05) RAGE (45 kDa) protein expression in diabetic (D) mice than in non-diabetic (ND) controls. **(C)** Methylglyoxal (MGO)-derived hydroimidazolone H1 (MG-H1) levels, measured using an MG-H1 ELISA kit and normalized w.r.t. total protein, were found to be significantly (*p<*0.01) higher in HG-treated HRECs than in NG-treated cells. **(D)** Representative Western blot bands and cumulative densitometric analysis of whole mouse retinas (n=6/group) revealed significantly (*p*<0.05) higher MG-H1 levels in diabetic (D) mice than in their non-diabetic (ND) counterparts. **(E)** RT-qPCR analysis of HRECs treated with NG, MN (osmotic control) or HG revealed a significant (∼2-fold; *p*<0.001) increase in ICAM-1 mRNA levels under HG conditions. **(F)** Representative Western blot bands and cumulative densitometric analysis of whole mouse retinas (n=6/group) revealed an ∼1.7-fold (*p<*0.01) higher ICAM-1 (85 kDa) protein expression in diabetic (D) mice than in non-diabetic (ND) controls. **(G, H)** HRECs were either left untreated (UT) or treated with increasing doses of MGO (0, 2.5, 10, 25, 100 μM) for 10d prior to Western blot analysis. Representative protein bands and cumulative densitometric analysis (n=6, including technical duplicates) indicate that RAGE (45 kDa) and ICAM-1 (85 kDa) protein levels were maximally and significantly (*p*<0.01) increased at the 10 μM dose. **(I)** Representative fluorescent images of adherent THP-1 monocytes and subsequent cell count (n≥6 images/condition) indicated a 4-fold (*p*<0.0001) increase in monocyte adhesion to 10 μM MGO-treated vs untreated (UT) HRECs. Scale bar: 200 μm. **(J, K, L)** HRECs were either left untreated (UT) or treated with increasing doses of MGO (0, 2.5, 10, 25, 100 μM) for 10d prior to assessment of LOX **(J)** mRNA levels (using RT-qPCR), **(K)** protein expression (using Western blot), and (**L)** activity (using Amplex Red**-**based assay), all of which peaked at the 10 μM dose. All mRNA and protein levels were normalized w.r.t. GAPDH while LOX activity measurements were compared with a hydrogen peroxide standard curve. *In vivo* data are plotted as mean ± SD and *in vitro* data as mean ± SEM, with *p*<0.05 considered as statistically significant.

To directly implicate RAGE in HG-induced LOX upregulation and associated retinal EC activation, we treated HRECs with MGO, a highly reactive AGE precursor that forms MG-H1 (AGE) adducts and thereby selectively binds to and upregulates RAGE (24). Our dose-dependent studies revealed that 10 μM MGO maximally increases the levels of MG-H1 (by 1.45-fold; *p*<0.0001; **Fig. S1A**), RAGE mRNA (by ∼2-fold; *p*<0.05; **Fig. S1B**) and protein (by 1.75-fold; *p*<0.01; **Fig. 1G**), and ICAM-1 protein (by 2.5-fold; *p*<0.01; **Fig. 1H**) in HRECs. Consistent with an increase in ICAM-1 expression, 10 μM MGO-treated HRECs exhibited ∼4-fold greater (*p*<0.001) monocyte adhesion than untreated ECs (**Fig. 1I**), thereby validating MGO as a potent inducer of RAGE-associated HREC activation.

Importantly, this trend in MGO-induced RAGE and ICAM-1 upregulation was mirrored by LOX expression. Specifically, treatment of HRECs with MGO produced a dose-dependent increase in LOX mRNA (*p*<0.0001; **Fig. 1J**), protein (*p*<0.001; **Fig. 1K**), and activity (*p*<0.01; **Fig. 1L**) that peaked at the same 10 μM dose that maximally increased RAGE expression (**Fig. 1G and S1B**).

### RAGE is required for diabetes-induced upregulation of retinal endothelial LOX

LOX expression increases under proinflammatory conditions (7,13). Since HG and MGO simultaneously upregulate LOX and proinflammatory RAGE (shown in Fig. 1) (5), we asked whether HG and MGO increase retinal endothelial LOX via RAGE. Indeed, addition of a RAGE-specific inhibitor FPS-ZM1 (25) to HG- and MGO-treated HRECs prevented the significant LOX upregulation seen in these conditions (**Fig. 2A, B**). Since the AGE/RAGE signaling activates the proinflammatory transcription factor NF-κB, we further asked whether RAGE increases LOX via NF-kB. As **Fig. 2C** shows, pharmacological inhibition of NF-κB in MGO-treated HRECs prevented the expected increase in LOX mRNA expression.

**Figure 2:**
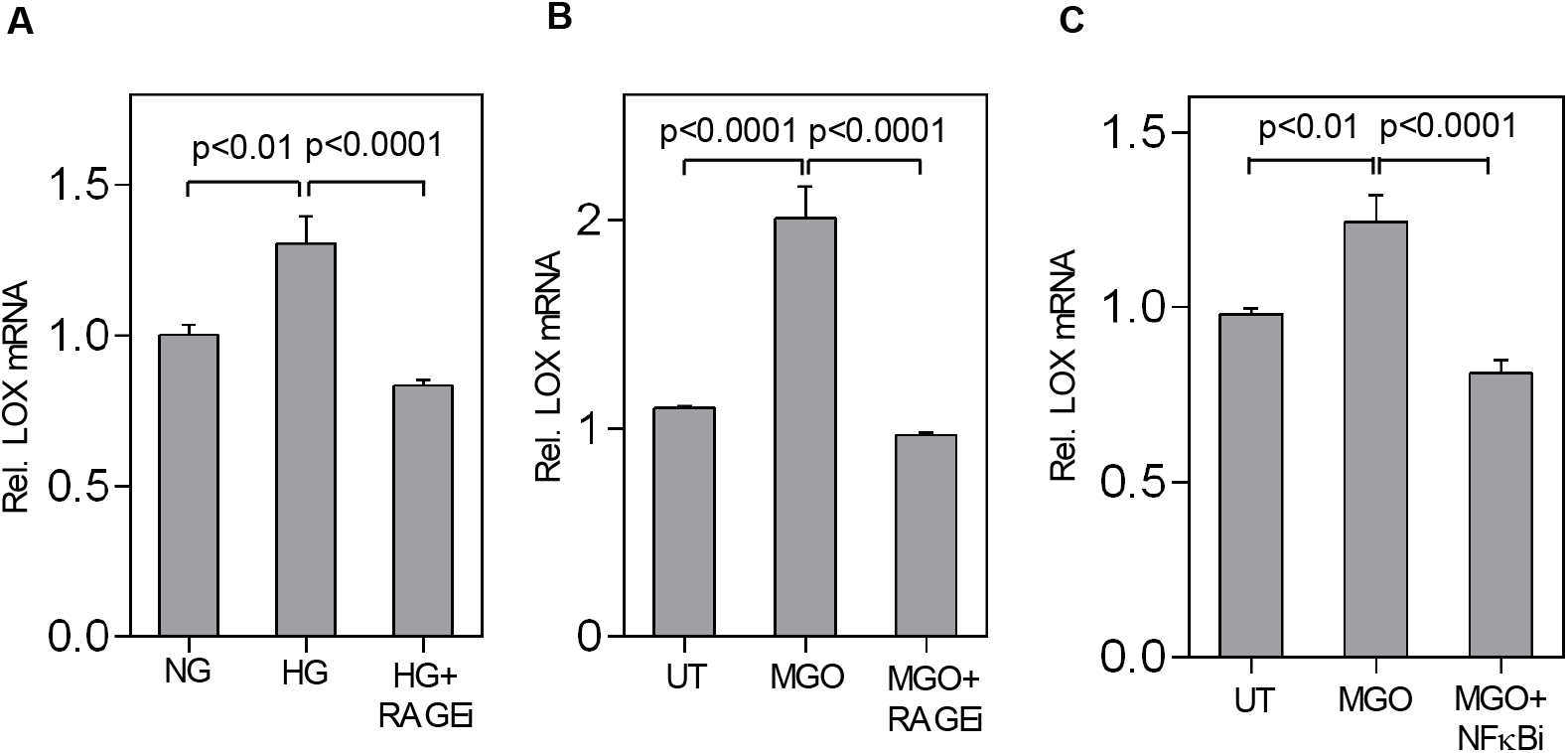
RAGE is required for diabetes-induced upregulation of retinal endothelial LOX. **(A, B)** RT-qPCR analysis of HRECs grown in medium containing NG, HG ± 0.1 μM RAGE inhibitor (RAGEi) FPS-ZM1 or MGO ± 0.1 μM RAGEi for 10d revealed that RAGEi completely blocks HG- or MGO-induced LOX mRNA upregulation (*p<*0.0001). **(C)** RT-qPCR analysis of HRECs that were either left untreated (UT) or treated with MGO ± NFκB inhibitor (NF-κBi) BAY 11-7082 (1 μM) for 3d showed that NF-κBi completely blocks MGO-induced LOX mRNA upregulation (*p*<0.0001). All mRNA and protein levels were normalized w.r.t. GAPDH. Data are plotted as mean ± SEM, with *p*<0.05 considered as statistically significant.

### LOX mediates MGO/RAGE-induced EC activation

Both MGO/RAGE **(Fig. 1)** and LOX (5) activate retinal ECs, leading to ICAM-1 upregulation. Since MGO/RAGE enhances LOX (shown in Fig. 1, 2), we asked whether MGO/RAGE promote retinal endothelial activation via LOX. Our findings revealed that simultaneously treating HRECs with MGO and BAPN, a pharmacological and irreversible inhibitor of LOX activity (26-28), prevents the MGO-induced increase in ICAM-1 mRNA **(Fig. S2)** and protein expression (**Fig. 3A**). To rule out any potential non-specific effects of BAPN, we alternatively reduced LOX levels in MGO-treated HRECs using siRNA **(Fig. S3)**. Our RT-qPCR measurements revealed that LOX-specific siRNA (but not scrambled siRNA) inhibits the MGO-induced increase in ICAM-1 mRNA by 75% (w.r.t. untreated HRECs; *p*<0.01; **Fig. 3B**).

**Figure 3:**
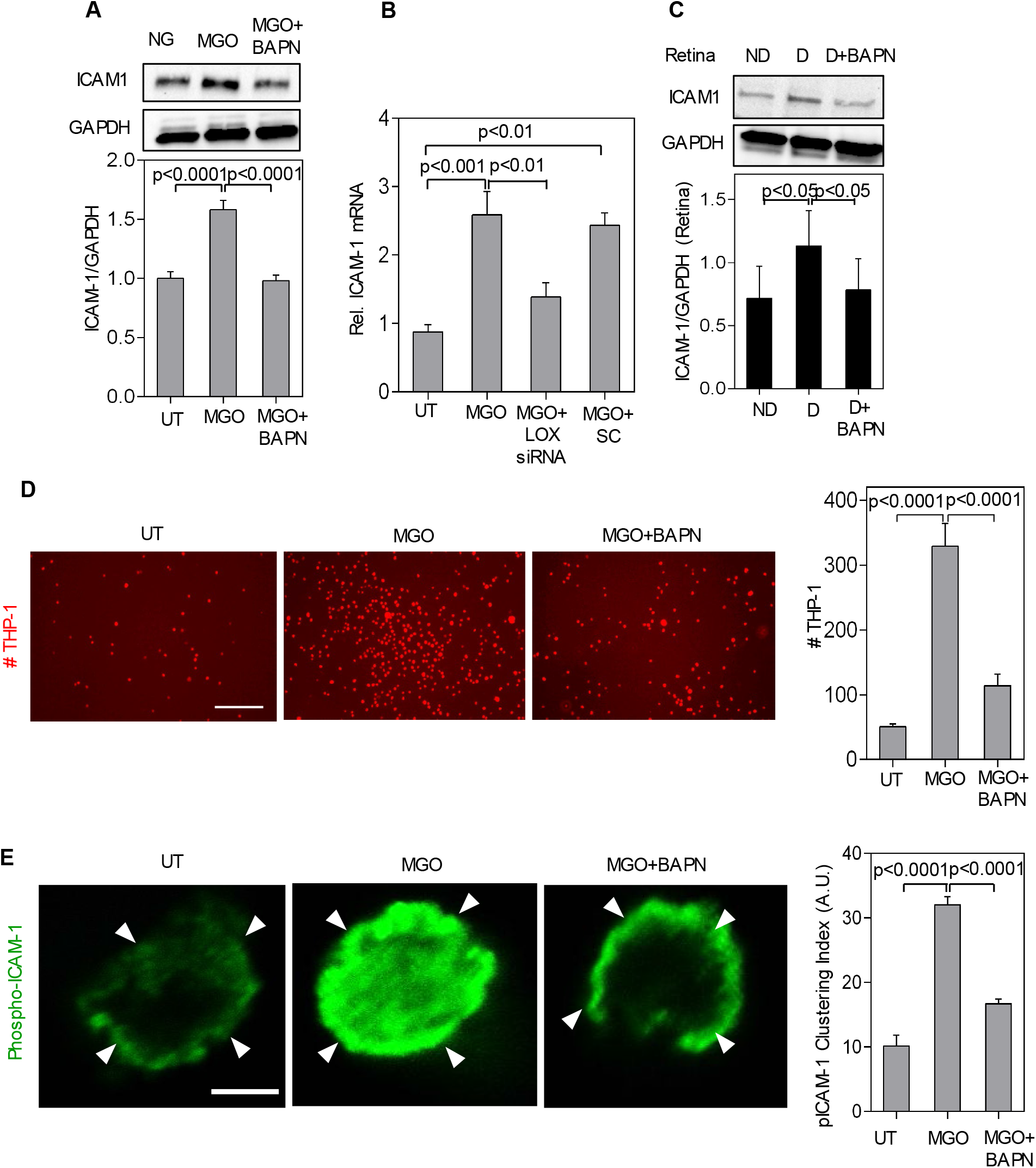
LOX mediates MGO/RAGE-induced EC activation. **(A)** HRECs were either left untreated (UT) or treated with MGO ± β-aminopropionitrile (BAPN; 0.1 mM) for 10d prior to Western blot analysis. Representative protein bands and cumulative densitometric analysis (n=6, including technical duplicates) indicate that LOX inhibition completely prevented the MGO-mediated ICAM-1 upregulation. **(B)** RT-qPCR analysis of HRECs that were either left untreated (UT) or treated with MGO ± LOX-specific or scrambled siRNA for 10d showed that silencing LOX reduces the MGO-induced ICAM-1 mRNA upregulation by 70% (*p*<0.01) and negative control scrambled siRNA did not show any effects on ICAM-1. **(C)** Representative Western blot bands and cumulative densitometric analysis of whole mouse retinas (n=6/group) showed that the 1.6-fold increase (*p*<0.05) in retinal ICAM-1 (85 kDa) seen in diabetic (D) mice is significantly suppressed (by 80%; *p*<0.05) by LOX inhibitor (3 mg/kg). **(D)** Representative fluorescent images of adherent THP-1 monocytes and subsequent cell count (n≥6 images/condition) revealed that the 6-fold (*p*<0.0001) increase in monocyte adhesion to MGO-treated HRECs was significantly inhibited by LOX inhibitor BAPN. UT: untreated HRECs. Scale bar, 400μm. **(E)** Immunolabeling of THP-1 monocyte-HREC co-cultures with anti-phospho-ICAM-1 and subsequent confocal imaging and intensity analysis revealed that MGO causes a 3-fold (*p*<0.0001) increase in ICAM-1 clustering (indicated by arrowheads) at monocyte-HREC adhesion site, which is significantly inhibited (by ∼75%; *p*<0.0001) by BAPN. Scale bar: 5 μm. All mRNA and protein levels were normalized w.r.t. GAPDH. *In vivo* data are plotted as mean ± SD and *in vitro* data as mean ± SEM, with *p*<0.05 considered as statistically significant.

Importantly, and consistent with these *in vitro* findings, LOX inhibition using BAPN almost completely blocked the retinal ICAM-1 upregulation seen in diabetic mice (**Fig. 3C**). Given this requirement of LOX in both MGO- and diabetes-induced increase in ICAM-1 expression in HRECs and mice retinas, respectively, BAPN predictably reduced (by 75% w.r.t. untreated HRECs; *p*<0.0001) the ICAM-1-dependent adhesion of THP-1 monocytes to MGO-treated HRECs (**Fig. 3D**). Interestingly, immunolabeling of the monocyte-EC adhesion sites with anti-phospho-ICAM-1, which labels activated ICAM-1, revealed that LOX inhibition also leads to a significant decrease (by 68% w.r.t. untreated HRECs; *p*<0.0001) in MGO-induced “clustering” of activated ICAM-1 around adherent monocytes (**Fig. 3E**). Thus, taken together, these findings establish LOX as a key mediator of the proinflammatory effects of MGO/RAGE on retinal ECs.

### LOX mediates MGO-induced subendothelial matrix stiffening

Both MGO-associated AGEs (29) and LOX (5) are implicated in the crosslinking and stiffening of subendothelial matrix. Since MGO enhances LOX (shown in Fig. 2), we asked whether MGO-induced subendothelial matrix crosslinking and stiffening are mediated by LOX. As LOX exists in both soluble and matrix-localized forms, we first decellularized the 15d HREC cultures (schematic; **Fig. S4**) and immunolabeled the residual subendothelial matrix with anti-LOX to confirm that MGO increases the levels of matrix-localized LOX. Fluorescence intensity analysis of LOX immunolabeled matrices revealed that MGO markedly increases (by 3-fold; *p*<0.001) matrix-localized LOX levels (**Fig. 4A)**, which was reduced by 70% (w.r.t. untreated HRECs; *p*<0.01) with BAPN treatment. This MGO-induced increase in matrix LOX caused a predictable increase (by 2.7-fold; *p*<0.0001) in subendothelial matrix stiffness that was nearly completely inhibited by BAPN **(Fig. 4B)**. Further, AFM imaging revealed that the normal ‘basketweave’ pattern of matrix fibers is modified by MGO treatment into a more compact and longitudinal alignment, which is notably reduced upon BAPN treatment **(Fig. 4C)**.

**Figure 4:**
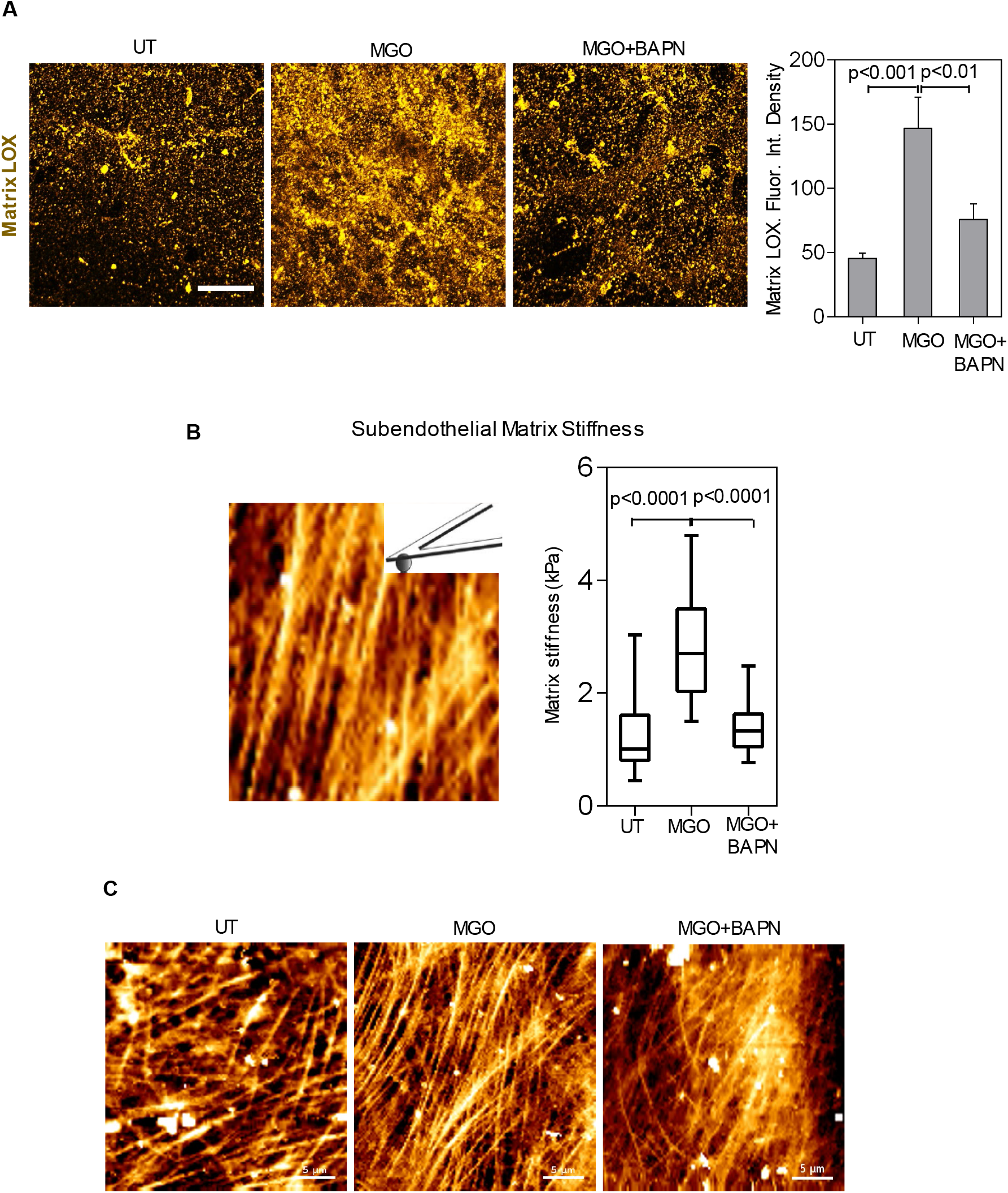
LOX mediates MGO-induced subendothelial matrix stiffening. **(A)** Decellularized matrices obtained from 15d-long untreated (UT) or MGO ± BAPN-treated HREC cultures were immunolabeled with anti-LOX. Representative confocal images and fluorescence intensity analysis (n≥6/condition) showed that MGO-treated HRECs deposit 3-fold higher amount (*p*<0.001) of matrix-localized LOX than untreated cells, which is significantly inhibited (by 70%; *p*<0.01) by BAPN treatment. Data are plotted as mean ± SEM. Scale bar, 20 μm. **(B)** Stiffness of unfixed decellularized subendothelial matrix was measured with a biological-grade atomic force microscope (AFM). Quantitative analysis of multiple (n≥75) force-indentation curves revealed a 1.7-fold increase (*p*<0.0001) in subendothelial matrix stiffness under MGO condition, which was significantly inhibited (by ∼80%; *p*<0.0001) by BAPN treatment. **(C)** Topographical scanning of subendothelial matrix using the AFM in QI^™^ mode reveal that MGO treatment modifies the normal basketweave pattern of matrix assembly into a longitudinal and compact organization, which appears to be prevented by BAPN. Scale bar, 5 μm. *p*<0.05 is considered as statistically significant.

### LOX mediates MGO/RAGE-induced EC activation through subendothelial matrix stiffening

We next asked whether matrix-localized LOX and associated matrix stiffening can mediate MGO/RAGE-induced EC activation independent of any potential effects of soluble LOX. To this end, we plated untreated HRECs on decellularized matrices obtained from preceding untreated or MGO ± BAPN-treated EC cultures and assessed ICAM-1 expression (schematic; **Fig. S4**). Under these conditions where elevated LOX is present only in the predeposited matrix, untreated HRECs plated on decellularized matrix from preceding MGO-treated cultures exhibited a substantial (*p*<0.01) increase in ICAM-1 mRNA expression, which was significantly inhibited (*p*<0.05) in cells plated on LOX-inhibited matrices **(Fig. 5A)**.

**Figure 5:**
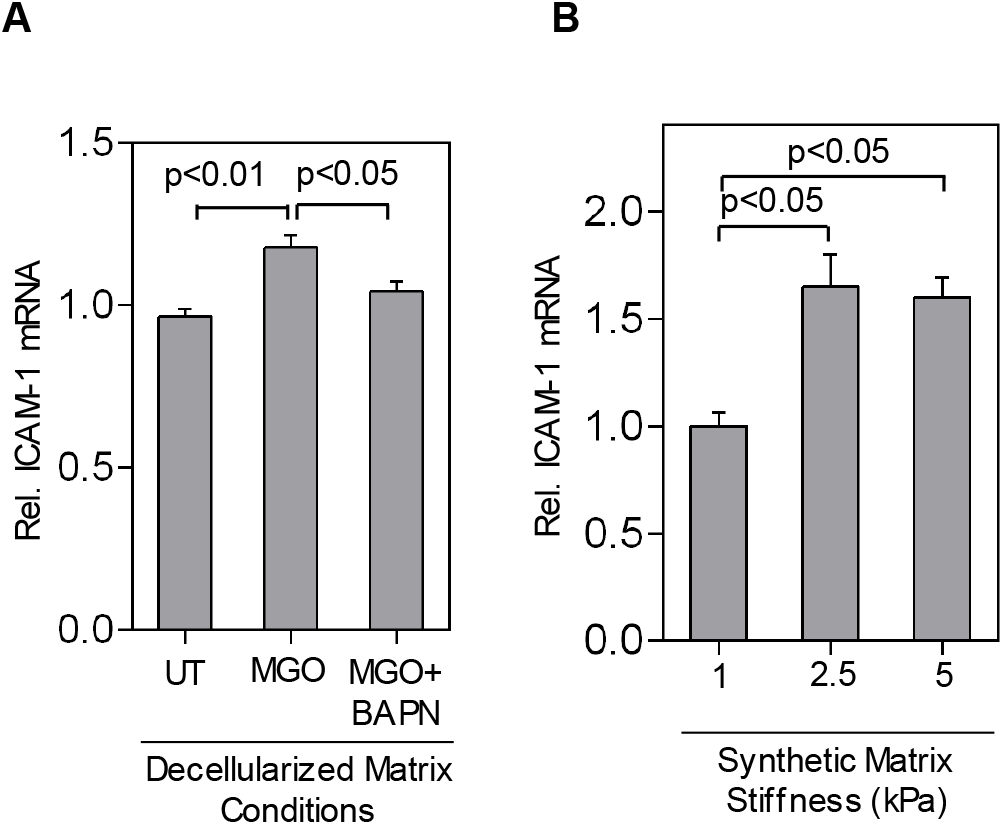
LOX mediates MGO/RAGE-induced EC activation through subendothelial matrix stiffening. **(A)** RT-qPCR analysis of HRECs plated on decellularized matrices obtained from preceding untreated (UT) or MGO±BAPN-treated HREC cultures show that the significant increase in ICAM-1 mRNA expression caused by MGO-treated matrix is inhibited on BAPN-normalized matrix. **(B)** RT-qPCR analysis of untreated (UT) HRECs plated on polyacrylamide-based synthetic matrices of increasing stiffness revealed that ICAM-1 mRNA expression increases significantly (by 1.6-fold; *p*<0.05) and peaks on the 2.5 kPa matrix. Data are plotted as mean ± SEM, with *p*<0.05 considered as statistically significant.

Since MGO can simultaneously alter matrix stiffness (shown in Fig. 4B) and density/composition (30), the aforementioned effect of MGO-treated matrix on fresh HRECs may result from changes in both mechanical (stiffness) and biochemical (ligand density/composition) cues from the matrix. To evaluate the independent effect of matrix stiffness on HREC activation, HRECs were plated on ‘synthetic’ matrices of tunable stiffness that were fabricated to mimic the subendothelial matrix stiffness of untreated (1 kPa) or MGO-treated (2.5 or 5 kPa) conditions (shown in Fig. 4B) and assessed for ICAM-1 expression. When compared with cells plated on the normal 1 kPa matrix, those plated on the stiffer 2.5 or 5 kPa matrices exhibited a 1.6-fold increase (*p*<0.05) in ICAM-1 mRNA levels **(Fig. 5B)**. Taken together, these data from decellularized and synthetic matrices indicate that the increase in subendothelial matrix stiffness is a crucial mediator of MGO/RAGE-induced HREC activation.

### LOX-dependent matrix stiffening feeds back to increase RAGE expression

Past studies have shown that inhibition of LOX activity reduces its own expression, thus indicating that LOX can auto-regulate its expression (31-33). Our findings agree with these observations as inhibiting LOX activity in HRECs with BAPN **(Fig. S5A)** completely blocked the MGO-induced increase in LOX mRNA **(Fig. S5B**) and protein **(Fig. S5C**) expression. Since RAGE promotes LOX expression (shown in Fig. 2), we wondered whether LOX auto-regulates its expression by feeding back to increase RAGE. Indeed, inhibiting LOX activity in HRECs with BAPN completely blocked the MGO-induced increase in RAGE mRNA (**Fig. 6A**) and protein (**Fig. 6B**) expression. Notably, siRNA-based silencing of LOX had the same inhibitory effect on RAGE expression, which was not seen with scrambled siRNA (**Fig. 6C**).

**Figure 6:**
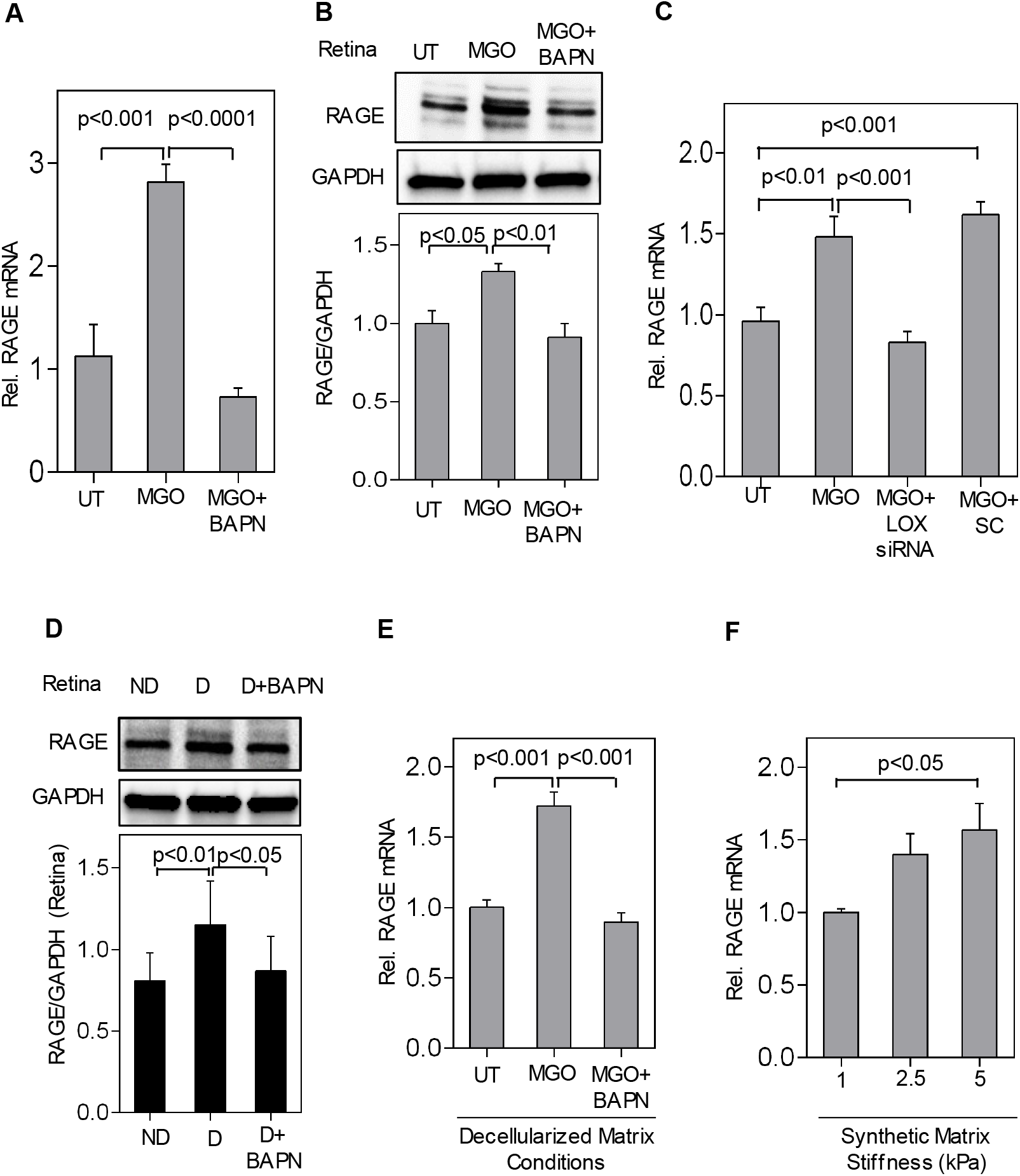
LOX-mediated matrix stiffening feeds back to increase RAGE expression. **(A, B)** RT-qPCR and Western blot analysis of HRECs treated with MGO ± BAPN for 10d revealed that LOX inhibition prevents MGO-induced RAGE mRNA and protein expression, respectively. **(C)** RT-qPCR analysis of HRECs transfected with LOX-specific or scrambled siRNA, followed by MGO treatment for 10d, shows that inhibiting LOX mRNA blocks MGO-induced RAGE mRNA upregulation. **(D)** Representative Western blot bands and cumulative densitometric analysis of whole mouse retinas (n=6/group) demonstrate that inhibiting LOX activity in diabetic (D) mice (using BAPN) prevents the diabetes-induced RAGE protein upregulation. ND: non-diabetic. **(E)** RT-qPCR analysis of HRECs plated on decellularized matrices obtained from preceding untreated (UT) or MGO±BAPN-treated HREC cultures show that the significant increase in RAGE mRNA expression caused by MGO-treated matrix is prevented on BAPN-normalized matrix. **(F)** RT-qPCR analysis of untreated (UT) HRECs plated on polyacrylamide-based synthetic matrices revealed that RAGE mRNA expression increases progressively with increasing matrix stiffness. Data are plotted as mean ± SEM, with *p*<0.05 considered as statistically significant.

The ability of LOX to feedback and promote RAGE expression was validated in vivo where diabetic mice treated with LOX inhibitor BAPN exhibited a near complete inhibition in retinal RAGE protein expression (**Fig. 6D**).

To determine whether LOX-dependent matrix stiffening is alone sufficient to promote autoregulation (via enhancement of RAGE expression) independent of any potential effects of soluble LOX, we plated fresh HRECs on decellularized matrices obtained from preceding untreated or MGO ± BAPN-treated EC cultures and assessed RAGE and LOX expression (schematic; **Fig. S4**). Our findings revealed that untreated HRECs plated on decellularized matrix from preceding MGO-treated cultures exhibit a significant increase in both RAGE (by 1.7-fold; *p*<0.001; **Fig. 6E**) and LOX (by 1.6-fold; *p*<0.001; **Fig. S6A**) mRNA levels, an effect that is completely blocked on LOX-inhibited matrices of normal stiffness.

To unequivocally confirm the role of subendothelial matrix stiffening in RAGE expression and, consequently, LOX autoregulation, HRECs were plated on the aforementioned synthetic matrices of normal (1 kPa) or increased (2.5 and 5 kPa) stiffness and assessed for RAGE and LOX mRNA. Similar to ICAM-1 expression reported in Fig. 5B, mRNA levels of both RAGE (**Fig. 6F**) and LOX **(Fig. S6B)** increased significantly (by ∼1.5-fold and ∼1.7-fold, respectively; *p*<0.05) on the stiffer matrices.

Taken together, these data from decellularized and synthetic matrices indicate that LOX-dependent subendothelial matrix stiffening alone can feed back to enhance RAGE expression and thereby achieve LOX autoregulation.

## Discussion

Despite the crucial role of retinal EC activation in inflammation-mediated DR pathogenesis (3,4,34), the underlying mechanisms remain poorly understood. We previously discovered that LOX-mediated subendothelial matrix stiffening contributes significantly to retinal EC activation under HG condition (5). The current study further extends our understanding of this mechanical control of retinal EC activation by shedding light on both the regulation and proinflammatory role of LOX. Specifically, we show that retinal endothelial LOX expression under hyperglycemic conditions is regulated by RAGE whose own expression, in turn, is enhanced by LOX-mediated subendothelial matrix stiffening. This sets in motion a mechanically regulated feed forward loop that sustains retinal EC activation in diabetes. Thus, these findings are significant and innovative as they provide new insights into retinal endothelial LOX regulation in diabetes, further highlight the role of mechanical cues in retinal EC activation, and implicate LOX as an alternative (downstream) target to block AGE/RAGE signaling that is otherwise difficult to achieve (17-19).

Vascular stiffening caused by an increase in LOX-mediated subendothelial matrix stiffness is being increasingly recognized as a key determinant of vascular inflammation associated with debilitating conditions such as atherosclerosis and sepsis (7,13). In line with these findings, we recently reported that LOX-mediated subendothelial matrix stiffening contributes significantly to HG-induced retinal EC activation (ICAM-1 expression) (5), a rate-limiting step in retinal vascular inflammation associated with DR (3,4,34). This finding has been supported by an independent study where LOX knockout mice did not develop the vascular lesions of DR (35). Although the exact mechanism of action was not investigated in this study, we speculate that this protective effect of LOX deficiency is due to the prevention of retinal vascular stiffening that, in turn, prevents retinal EC activation. Despite these strong implications of retinal endothelial LOX in DR, a mechanistic understanding of its regulation and proinflammatory role in diabetes remains absent.

Past and current findings reveal that, under HG/diabetic conditions, LOX-mediated retinal EC activation is associated with upregulation of proinflammatory RAGE, which promotes NF-kB activation and vascular inflammation (5,16,36). LOX-mediated vascular inflammation in other inflammatory conditions such as sepsis and atherosclerosis are also associated with RAGE overexpression (37,38). Thus, we asked whether RAGE could directly increase retinal endothelial LOX. Although RAGE is a multiligand receptor, its primary ligand are the AGEs that result from nonenzymatic glycation of proteins and lipids caused by HG and the highly reactive glycolysis metabolites such as MGO. Indeed, MG-H1, an MGO-derived AGE, is abundant in the retinas of diabetic rodents (36,39), which is consistent with a significantly higher MG-H1 levels we see in both HG-treated HRECs and retinas of diabetic C57BL/6J mice. These findings, coupled with past reports of elevated MGO levels in diabetic patients (40,41), led us to choose the AGE precursor MGO as the RAGE ligand for the current study.

We found that MGO treatment for 10d maximally increases HREC LOX expression at the same dose that causes peak RAGE expression and HREC activation (ICAM-1 expression). Subsequent studies confirmed that this MGO-dependent increase in LOX levels is mediated by RAGE because treating HRECs with a RAGE inhibitor blocked the MGO-induced LOX upregulation. This is, to our knowledge, the first direct evidence of LOX upregulation by MGO and RAGE and aligns with previous findings that NF-κB, a downstream effector of RAGE, can increase LOX transcription in ECs and cancer cells (42,43). Interestingly, contrary to these findings, a separate study found LOX expression to be unchanged in non-differentiated mouse osteoblasts treated with AGE-modified collagen for 24h. Thus, it appears that the LOX-enhancing effect of AGEs or their precursors is dependent on the nature of the treated cells, their degree of differentiation, and/or the treatment period. Nonetheless, given our finding that RAGE enhances retinal endothelial LOX, it is possible that other proinflammatory cytokine receptors, which are simultaneously upregulated in DR and cause NF-kB activation, also increase retinal endothelial LOX levels. Addressing this possibility will be the focus of our future studies.

We have previously shown that LOX promotes retinal EC activation under HG conditions (5). Our current findings indicate that LOX achieves this effect, in large part, by mediating the proinflammatory effects of RAGE because LOX inhibition nearly completely blocked the MGO-induced HREC activation (ICAM-1 expression and monocyte-EC adhesion) in vitro and diabetes-induced retinal ICAM-1 expression in vivo. Independent studies have shown that LOX inhibition blocks the proinflammatory effects of lipopolysaccharide (LPS) on pulmonary ECs (7,44) and ApoE knockout on aortic ECs (13). LOX activity is also significantly increased in aortas of atheroprone animals (45). Thus, LOX appears to play a central role in vascular inflammation caused by multiple risk factors. Interestingly, LOX inhibition also blocked the MGO-induced ICAM-1 ‘clustering’ at the site of monocyte-HREC adhesion. We have previously shown that endothelial ICAM-1 clustering around adherent monocytes can be induced by subendothelial matrix stiffening (12). Consistent with this finding, our AFM measurements show that LOX inhibition also significantly reduced MGO-induced subendothelial matrix stiffening. That LOX inhibition did not return subendothelial matrix stiffness to the basal (untreated) level indicates that MGO also directly crosslinks and stiffens matrix by forming AGE adducts (46). Overall, these findings demonstrate that LOX is a crucial mediator of both the biochemical (retinal EC activation) and mechanical (subendothelial stiffening) effects of MGO/RAGE.

LOX exists in both matrix-localized and soluble forms. While soluble LOX has primarily been implicated in cell migration (47,48), the aforementioned proinflammatory effects of LOX have been attributed to its matrix-localized form where it promotes subendothelial matrix stiffening and, thereby, alters EC mechanotransduction (5,7,13,44). Consistent with the latter reports, here we found that MGO-induced LOX upregulation is also significantly localized to the subendothelial matrix where it causes matrix stiffening. To unequivocally confirm the role of LOX-mediated subendothelial matrix stiffening in MGO-induced retinal EC activation, we cultured HRECs on decellularized matrices obtained from long-term HREC cultures or on polyacrylamide-based synthetic matrices of tunable stiffness that confer independent control of matrix stiffness vs composition (5,12). Findings from these studies revealed that subendothelial matrix stiffening plays a crucial role in promoting MGO-induced HREC activation. We had previously shown that HG-induced HREC activation is also similarly dependent on LOX-mediated subendothelial matrix stiffening (5). Thus, taken together, these findings underscore the pivotal role that LOX plays in the matrix-mediated mechanical control of retinal EC activation associated with key DR risk factors.

Although these findings reveal the RAGE-mediated regulation and proinflammatory role of retinal endothelial LOX in diabetes, they do not explain precisely how LOX promotes retinal EC activation. In this regard, we interestingly found that inhibiting LOX activity inhibits its own mRNA expression in MGO-treated HRECs and retinas of diabetic mice, thereby indicating that LOX can autoregulate its expression. This finding is consistent with previous reports of LOX autoregulation in animal models of tissue fibrosis (31,33). Notably, in contrast to transcription factors, proteins/enzymes such as LOX require the involvement of additional factor(s) for their autoregulation (49). As we found out, that additional factor for LOX autoregulation is RAGE that, we already showed, enhances LOX expression. Specifically, we found that LOX autoregulates its expression by feeding back to increase RAGE expression because LOX inhibition completely blocked RAGE upregulation in both MGO-treated HRECs and retinas of diabetic mice. We further show that this LOX-RAGE feedback signaling is driven primarily by LOX-dependent subendothelial matrix stiffening. Taken together, these findings indicate that RAGE-regulated retinal endothelial LOX exerts proinflammatory effects in DR by increasing subendothelial matrix stiffness that, in turn, feeds back to control RAGE expression (**Figure 7-Schematic**). Whether LOX can similarly (mechanically) control the expression of other DR-associated proinflammatory factors such as cytokines, their receptors, and intracellular signaling complexes remains to be seen.

**Figure 7:**
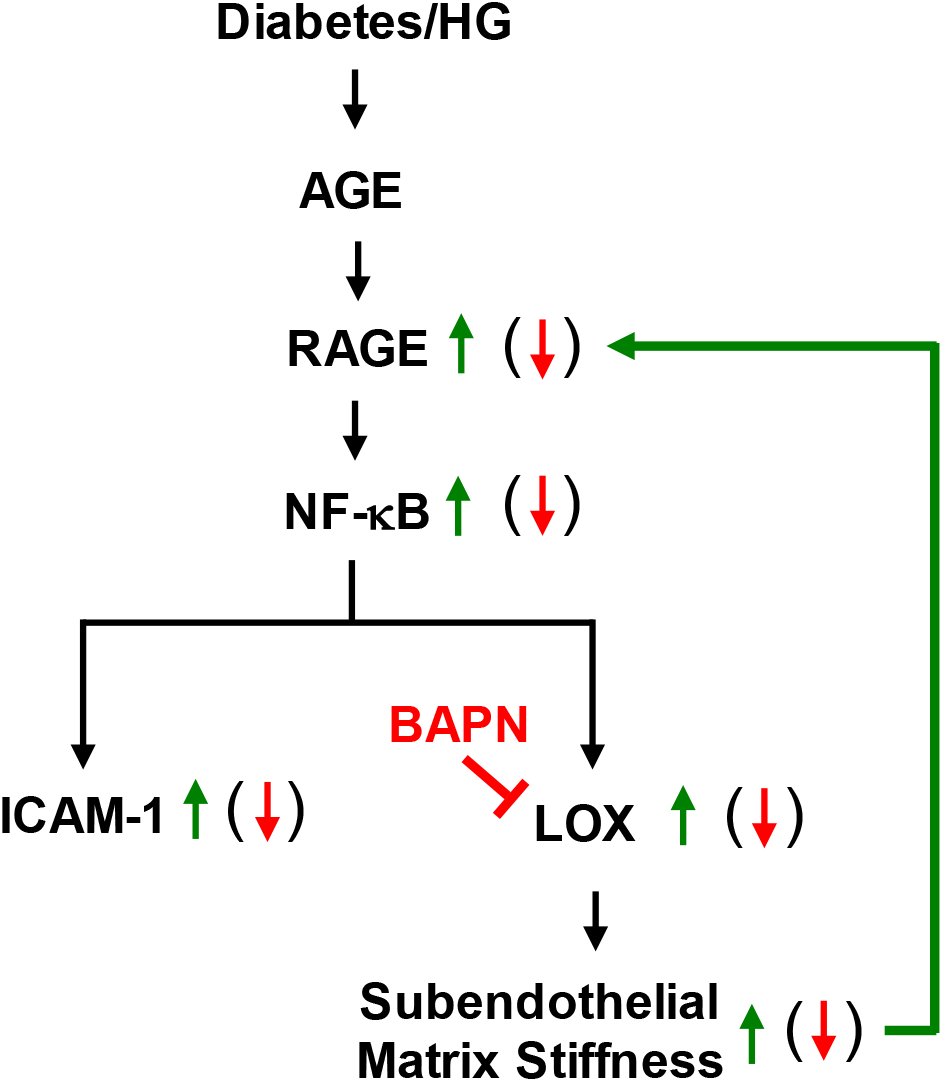
Schematic- A potential mechanism to explain the LOX-dependent mechanical control of retinal endothelial RAGE. Based on our current findings, we propose that LOX establishes the mechanical control of RAGE-dependent retinal EC activation and vascular inflammation in DR. This newly identified mechanism of DR pathogenesis begins with the formation of AGEs, the key RAGE ligands that accumulate and are implicated in DR. AGE/RAGE interaction triggers downstream activation of the master inflammatory transcriptional factor NF-kB that simultaneously promotes ICAM-1 and LOX expression in retinal ECs. While ICAM-1 expression (EC activation) directly leads to vascular inflammation, upregulation of LOX results in subendothelial matrix remodeling in the form of increased matrix stiffness. This stiffer matrix, in turn, feeds back to further enhance retinal endothelial RAGE expression, and consequently retinal EC activation and vascular inflammation, via a yet unknown mechanotransduction pathway.

Our finding that LOX-dependent subendothelial matrix stiffening both regulates and is regulated by RAGE is significant because it offers fresh mechanistic insights into the regulation and proinflammatory role of LOX in diabetes. Perhaps more crucially, the ability of LOX to regulate RAGE has important translational implications because LOX might serve as an alternative (downstream) target to block AGE/RAGE signaling in retinal ECs, thus overcoming the limitations of direct AGE/RAGE targeting in DR such as drug toxicity and low efficacy (17-19). If so, it would mark a significant advancement in the future clinical management of DR.

## Author Contributions

S.C. designed and performed experiments, analyzed data, and wrote the manuscript, I.S.T., M.A., and N.M. performed experiments and analyzed data, T.S.K. designed experiments, and K.G. conceived the idea, designed experiments, analyzed data, and wrote the manuscript. All authors reviewed, edited, and approved the manuscript. K.G. is the guarantor of this work and, as such, takes responsibility for the integrity and accuracy of the reported data.

