## Supplemental Material for "Lysyl oxidase-dependent subendothelial matrix stiffening promotes RAGE-mediated retinal endothelial activation in diabetes"

#### Research Design and Methods

##### Experimental Animals

All animal procedures were performed in accordance with the Association for Research in Vision and Ophthalmology (ARVO) Statement for the Use of Animals in Ophthalmic and Vision Research and approved by University of California Riverside Institutional Animal Care and Use Committee. Diabetes was induced in adult (8 wk-old) male C57BL/6J mice (Jackson Laboratory, Bar Harbor, ME, USA) through daily i.p. injections of freshly-prepared streptozotocin solution (STZ; MP Biomedicals, Irvine, CA; 60 mg/kg body weight in 10 mM citrate buffer; pH 4.5) for five consecutive days. Mice with fasting blood glucose >275 mg/dL two weeks after the last STZ injection were classified as diabetic, with this time point being considered as the onset of overt diabetes. Age-matched normal C57bL/6J mice receiving citrate buffer alone served as non-diabetic controls.

##### Cell culture

Human THP-1 monocytes were purchased from ATCC (Manassas, VA, USA) and cultured in RPMI medium (catalog no.SH30096.01; Hyclone) supplemented with 10% fetal bovine serum (FBS), 1X GlutaMAX, and Penicillin-Streptomycin (5000 Units/mL; catalog no.15070063; Gibco™-Thermo Fisher Scientific), 1 mM sodium pyruvate (catalog no.11360070; Gibco™-Thermo Fisher Scientific) and 10 mM HEPES (catalog no.11360-070; Thermo Fisher Scientific-Gibco).

##### RT-qPCR

Total RNA was isolated from HRECs (Direct-zol RNA MiniPrep; Zymo Research, Irvine, CA, USA), converted to cDNA using a High-Capacity RNA-to-cDNA kit (catalog no. 4387406;

Thermo Fisher Scientific-Applied Biosystems™), and amplified with gene/species-specific TaqMan primers for RAGE (Hs00542584_g1), LOX (Hs00942480_m1) and ICAM-1 (Hs00164932_m1) in QuantStudio™ 5 Real-Time PCR system. Target gene expression was normalized to the house keeping gene GAPDH (Hs02786624_g1) and relative expression was determined using the comparative Delta Delta Ct (DDCt) method. Negative control was performed for each reaction assay plates.

**Western blot**

To determine the expression levels of target proteins, equal amounts (20 or 40 μg) of total protein were separated in 4-15% Mini-PROTEAN® TGX precast protein gels (Biorad, Hercules, CA, USA) and transferred to nitrocellulose membrane prior to probing with primary antibodies against RAGE (catalog no. ab3611; Abcam), MG-H1 (catalog no. STA-011; Cell Biolabs, Inc., San Diego, CA, USA), LOX (catalog no. NB110-59729; Novus Biologicals, San Diego, CA, USA), ICAM-1 (catalog no. SC-8439; Santa Cruz Biotechnology, Santa Cruz, CA, USA), and loading control GAPDH (catalog no. G9545; Sigma-Aldrich), followed by appropriate secondary antibody conjugated to horseradish peroxidase (Catalog no. PI-1000 and PI-2000; Vector Laboratories). Protein bands were detected using a SuperSignal™ West Dura Extended Duration Substrate (Catalog no. 34075; Thermo Fisher Scientific, Waltham, MA, USA) and imaged using ChemiDoc XRS+ System (Biorad). Each target was assessed from technical duplicates obtained from three independent experiments. ImageJ was used to perform densitometric analysis.

**Monocyte-EC adhesion assay**

Monocyte-EC adhesion assay was performed as per our previously reported protocol (1). Briefly, 10 day-long HREC cultures were serum starved (2.5% fetal bovine serum) overnight prior to addition of DAPI-labeled THP-1 monocytes (passage 5-10) at 125,000 cells/cm$^2$ for 30 min at

37°C. After washing away the non-adherent monocytes with PBS, the co-culture was fixed with 1% PFA and imaged with an epifluorescence Nikon Eclipse TS2 microscope for monocyte counting using ImageJ (n≥6 images per condition). Co-cultures were subsequently labeled with rabbit anti-phospho ICAM-1 (Tyr512, GeneTex, Irvine, CA, USA) followed by addition of fluorescently-labeled anti-rabbit IgG (Jackson ImmunoResearch Laboratories, Inc.) to visualize phospho ICAM-1 clustering using Zeiss LSM 710 confocal microscope (Zeiss). ICAM-1 clustering index was quantified by measuring phosphor ICAM-1 fluorescence intensity (n≥6 images per condition) at the monocyte-EC adhesion site and normalizing it to the average 'background' intensity measured from three neighboring cytoplasmic sites, as we have reported before (2).

**LOX Activity assay**

LOX activity was measured from HREC culture supernatant using a conventional Amplex™ Red fluorescence assay, as reported previously (1, 3). The fluorescence signal, resulting from hydrogen peroxide produced by active LOX, was measured at 540/590 nm in SpectraMax iD5 multimode plate reader (Molecular Device) and the LOX activity was determined by comparing intensity values with a standard curve generated from serially-diluted hydrogen peroxide solution.

**Subendothelial matrix**

Decellularized subendothelial matrix was obtained using our previously reported protocol (4). Briefly, HRECs in NG or MGO ± BAPN medium were grown on activated glass coverslips for 15d prior to decellularization of the cell monolayers with mild detergent (composed of 20 mM ammonium hydroxide and 0.5% Triton X-100) that removes the cells without hampering the EC-secreted matrix. The decellularized matrices were subsequently treated with DNase to remove cell debris before they were either fixed using 0.5% PFA for 15 min at 4°C for immunofluorescence staining or left unfixed (fresh) for matrix stiffness/topography measurement and cell replating assays. To visualize matrix-localized LOX, fixed decellularized matrices were labeled with rabbit anti-LOX antibody (Novus Biologicals) followed by addition of a fluorescently-labeled anti-rabbit IgG (Jackson ImmunoResearch Laboratories, Inc.), and imaged using Zeiss LSM 710 confocal microscope. ImageJ was used to quantify LOX fluorescence intensity ($n \geq 6$ images per condition) and represented as integrated density.

**Synthetic matrix fabrication**

The synthetic matrices used in the current study were fabricated as per the previously published protocols (1, 2, 5). Briefly, thin (~100 µm thick) elastic polyacrylamide gels of 1000 Pascals (Pa), 2500 Pa, and 5000 Pa were fabricated to mimic subendothelial matrix stiffness in normal or diabetic conditions and coated with the proinflammatory matrix molecule fibronectin (5 $\mu g/cm^2$). HRECs were next plated on these synthetic matrices in regular culture medium for 24 h prior to mRNA isolation for RT-qPCR analysis.

#### Supplementary Figure Legends

**Figure S1: MGO dose dependently increases MG-H1 and RAGE levels in HRECs**

**(A)** MG-H1 levels were measured in HREC lysates using an ELISA kit following treatment with increasing doses of MGO (2.5, 10, 25, 100 µM) for 10 d. Results indicate that MG-H1 level, normalized w.r.t. protein concentration, was maximally ($p$<0.0001) increased at the 10 µM dose. **(B)** RT-qPCR analysis of HRECs treated with increasing doses of MGO for 10 d revealed the highest increase in RAGE mRNA ($p$< 0.05) at the 10 µM dose. Data are plotted as mean ± SEM, with $p$<0.05 considered as statistically significant.

**Figure S2: LOX inhibition prevents MGO-induced increase in endothelial ICAM-1**

RT-qPCR analysis of HRECs treated with MGO ± BAPN for 10 d reveal that LOX inhibition by BAPN significantly prevents MGO-induced ICAM-1 mRNA levels ($p$< 0.01).

**Figure S3: LOX-specific siRNA blocks LOX expression in HRECs**

RT-qPCR analysis of LOX mRNA levels in HREC cultures transfected with LOX or scrambled siRNA followed by MGO treatment for 10d shows that LOX-specific siRNA completely blocks MGO-induced LOX mRNA upregulation while scrambled siRNA (negative control) showed no effects. Data are plotted as mean ± SEM, with $p$<0.05 considered as statistically significant.

**Figure S4: Schematic illustration of the process to obtain decellularized matrix**

The schematic depicts the experimental procedure to obtain decellularized subendothelial matrix prior to measurement of matrix protein expression (using immunostaining), matrix stiffness (using AFM), or matrix-dependent HREC activation (by replating fresh HRECs on decellularized matrices).

**Figure S5: Inhibiting LOX activity inhibits its own expression**

**(A)** Amplex™ Red fluorescence assay performed on HRECs treated with MGO±BAPN for 10d shows BAPN prevents MGO-induced increase in LOX activity ($p$<0.01). **(B, C)** RT-qPCR and Western Blot analysis of HRECs treated with MGO±BAPN for 10 d revealed that inhibiting LOX activity using BAPN prevents MGO-induced upregulation of both LOX mRNA and protein (32 kDa) expression ($p$<0.001). Data are plotted as mean ± SEM, with $p$<0.05 considered as statistically significant.

**Figure S6: Matrix stiffening increases LOX expression**

**(A)** RT-qPCR analysis of HRECs plated on decellularized matrices obtained from preceding untreated (UT) or MGO±BAPN-treated HREC cultures show that the significant increase in LOX mRNA expression caused by MGO-treated matrix is prevented on BAPN-normalized matrix. **(B)** RT-qPCR analysis of untreated (UT) HRECs plated on polyacrylamide-based synthetic matrices revealed that LOX mRNA expression increases progressively with increasing matrix stiffness. Data are plotted as mean ± SEM, with $p<0.05$ considered as statistically significant.

**Supplementary Figure 1**

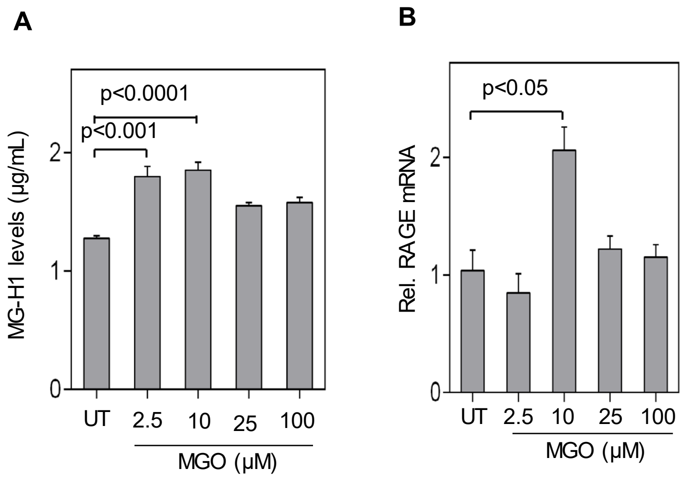

**Supplementary Figure 2**

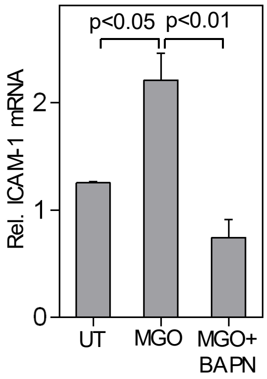

### Supplementary Figure 3

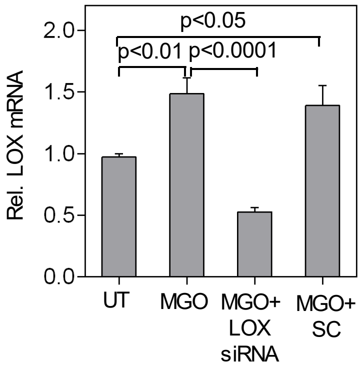

### Supplementary Figure 4

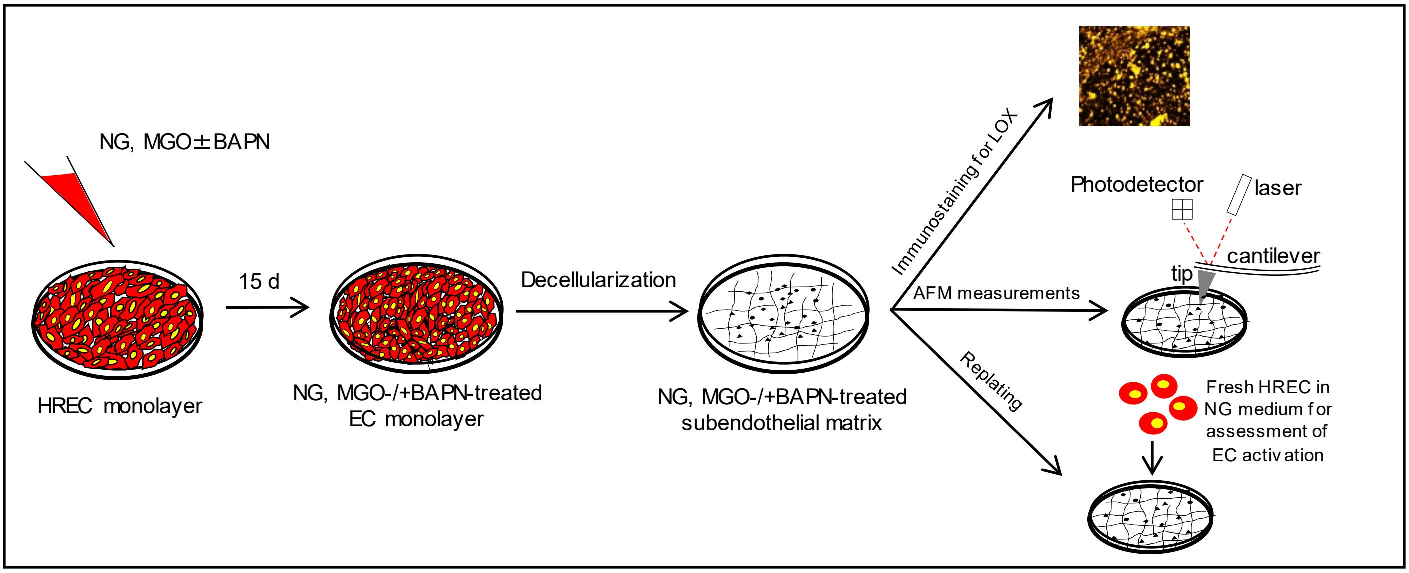

**Supplementary Figure 5**

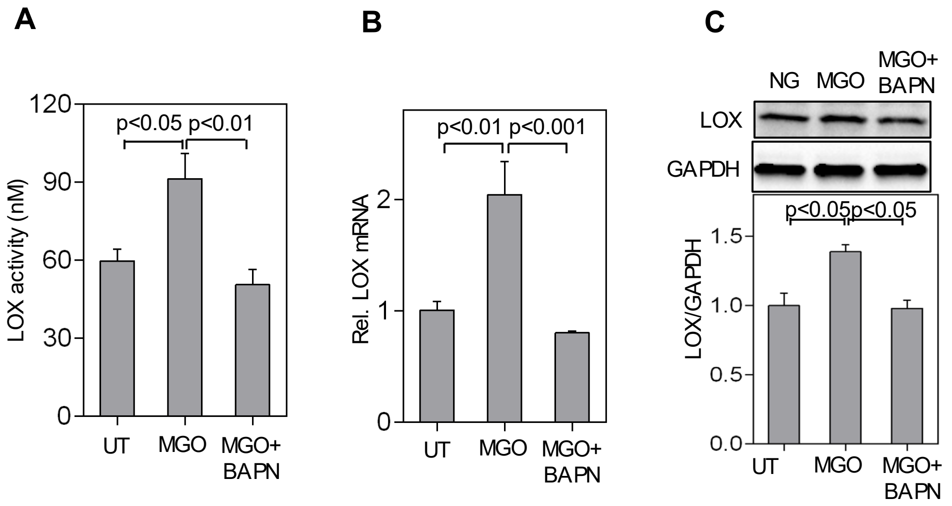

**Supplementary Figure 6**

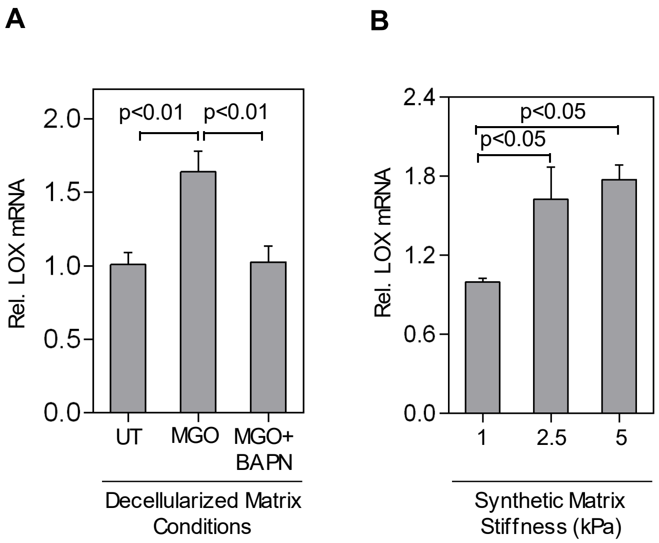
